# Neutralization and Stability of JN.1-derived LB.1, KP.2.3, KP.3 and KP.3.1.1 Subvariants

**DOI:** 10.1101/2024.09.04.611219

**Authors:** Pei Li, Julia N. Faraone, Cheng Chih Hsu, Michelle Chamblee, Yajie Liu, Yi-Min Zheng, Yan Xu, Claire Carlin, Jeffrey C. Horowitz, Rama K. Mallampalli, Linda J. Saif, Eugene M. Oltz, Daniel Jones, Jianrong Li, Richard J. Gumina, Joseph S. Bednash, Kai Xu, Shan-Lu Liu

## Abstract

During the summer of 2024, COVID-19 cases surged globally, driven by variants derived from JN.1 subvariants of SARS-CoV-2 that feature new mutations, particularly in the N-terminal domain (NTD) of the spike protein. In this study, we report on the neutralizing antibody (nAb) escape, infectivity, fusion, and stability of these subvariants—LB.1, KP.2.3, KP.3, and KP.3.1.1. Our findings demonstrate that all of these subvariants are highly evasive of nAbs elicited by the bivalent mRNA vaccine, the XBB.1.5 monovalent mumps virus-based vaccine, or from infections during the BA.2.86/JN.1 wave. This reduction in nAb titers is primarily driven by a single serine deletion (DelS31) in the NTD of the spike, leading to a distinct antigenic profile compared to the parental JN.1 and other variants. We also found that the DelS31 mutation decreases pseudovirus infectivity in CaLu-3 cells, which correlates with impaired cell-cell fusion. Additionally, the spike protein of DelS31 variants appears more conformationally stable, as indicated by reduced S1 shedding both with and without stimulation by soluble ACE2, and increased resistance to elevated temperatures. Molecular modeling suggests that the DelS31 mutation induces a conformational change that stabilizes the NTD and strengthens the NTD-Receptor-Binding Domain (RBD) interaction, thus favoring the down conformation of RBD and reducing accessibility to both the ACE2 receptor and certain nAbs. Additionally, the DelS31 mutation introduces an N-linked glycan modification at N30, which shields the underlying NTD region from antibody recognition. Our data highlight the critical role of NTD mutations in the spike protein for nAb evasion, stability, and viral infectivity, and suggest consideration of updating COVID-19 vaccines with antigens containing DelS31.

## INTRODUCTION

A global surge in COVID-19 cases has been ongoing since the beginning of summer 2024 and continues to rise. This year has been dominated by the circulation of the BA.2.86-derived JN.1 variant of SARS-CoV-2 and its descendants^1,2^. These variants are characterized by marked immune escape, making vaccinated and convalescent sera less effective, although immunity is somewhat improved with the most recent XBB.1.5 spike mRNA monovalent vaccine formulation or repeated exposure to Omicron variants^3–12^. The JN.1 lineage of SARS-CoV-2 is continuing to accumulate mutations, showing distinct convergent evolution at key spike protein residues, including R346, F456, and, most recently, DelS31^1,2,13^. Currently, several of these variants are increasing in circulation, though the underlying mechanisms remain to be fully understood.

Throughout 2024, various JN.1-derived variants have fluctuated in prevalence. Early in the year, the JN.1 variant, characterized by a single L455S mutation relative to the parental BA.2.86 variant, was dominant^3,4^. This single mutation significantly enhanced the virus’s immune evasion and transmission^3,6,14,15^. However, variants like FLiRT, SLiP and KP.2 quickly supplanted JN.1, driven by key spike mutations R346T and F456L, which further contributed to immune evasion^8,9,16–21^. More recent variants are now developing mutations in other regions, particularly concentrated in the N-terminal domain (NTD) of the spike protein (**Fig 1A**). Globally, variants such as KP.2, KP.3, LB.1, KP.2.3, and KP.3.1.1 are on the rise (**Fig. 1B-C**)^1,2,13^, and a new deletion of residue S31 (DelS31) has emerged convergently in these variants^2^, suggesting the latter may play a role in viral fitness, though its exact consequences are still unknown.

**Figure 1:**
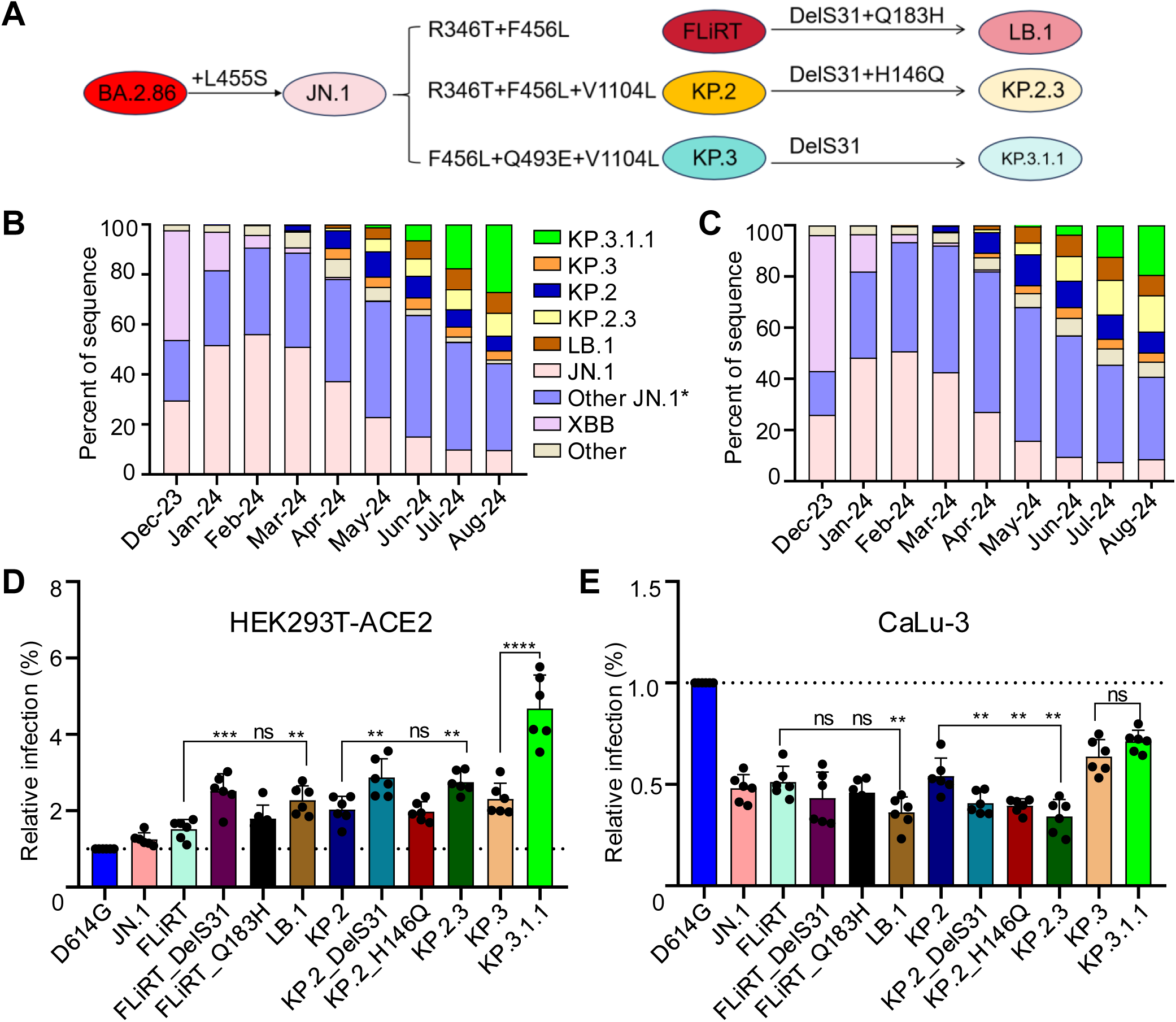
Infectivity of JN.1 subvariants in 293T-ACE2 and CaLu-3 cells. **(A)** Mutations that characterize JN.1-derived subvariants FLiRT, KP.2, KP.3, LB.1, KP.2.3, and KP.3.1.1. Recorded incidences of infection by relevant variants in **(B)** the United States and **(C)** globally based on data collected by the Centers for Disease Control and Prevention (CDC) and Global Initiative of Sharing All Influenza Data (GISAID). Other JN.1*: JN.1 subvariants. Infectivity of pseudotyped lentiviral vectors bearing variant spikes of interest was determined in **(D)** 293T-ACE2 cells and **(E)** CaLu-3 cells. Bars in **(D and E)** represent means and standard deviation from 6 independent infections (n=6). Significance was determined and displayed relative to D614G, stars represent ** p < 0.01; ***p < 0.001 and ns p > 0.05.

In this study, we focus on the variants currently dominating circulation in the United States, including LB.1, KP.2.3, KP.3, and KP.3.1.1. We investigated these variants in comparison to parental strains D614G, JN.1, FLiRT, and KP.2, as well as the impact of single mutations in the NTD, such as DelS31, H146Q, and Q183H. The goal of our study was to better understand the impact these new JN.1-derived variants have on nAb titers in individuals who received the bivalent mRNA vaccine, patients hospitalized during the BA.2.86/JN.1 wave in Columbus, Ohio, and a cohort of hamsters that received two doses of the monovalent XBB.1.5 vaccine. We also sought to understand the underlying mechanism by characterizing key aspects of spike biology, including infectivity, cell-cell fusion, processing, and stability.

## RESULTS

### Infectivity of LB.1, KP.2.3 and KP.3.1.1 in 293T-ACE2 cells and CaLu-3 cells

We first investigated the infectivity of lentivirus pseudotypes bearing SARS-CoV-2 spikes of recently emerged JN.1 subvariants in 293T cells overexpressing human ACE2 (293T-ACE2) (**Fig 1D**) and in the human lung-derived cell line CaLu-3 (**Fig 1E**). As we have established previously^9^, JN.1 and its derived FLiRT subvariants exhibited a modestly increased infectivity in 293T-ACE2 cells compared to ancestral D614G. Notably, the newly emerged KP.3 and KP.3.1.1 subvariants exhibited higher titers than parental JN.1. In particular, the addition of the DelS31 mutation in FLiRT-DelS31, KP.2-DelS31 and KP.3-DelS31 (i.e., KP.3.1.1) caused a 1.7-fold, 1.4-fold and 2.0-fold increase in titer relative to the parental FLiRT (p < 0.001), KP.2 (p < 0.01) and KP.3 (p < 0.001), respectively (**Fig 1D)**. Single point mutations Q183H in FLiRT (FLiRT_Q183H) and H146Q in KP.2 (KP.2_H146Q) did not appear to contribute to the increased infectivity of LB.1 (harboring both DelS31 and Q183H) and KP.2.3 (containing both DelS31 and H146Q) relative to their ancestral FLiRT and KP.2, respectively (**Fig 1D**).

In CaLu-3 cells, we observed distinct phenotypes. While all JN.1-derived variants exhibited much lower titers compared to ancestral D614G (**Fig. 1E**, p < 0.0001) as we have established previously^9,14^, KP.3.1.1 showed increased titers compared to parental JN.1. Interestingly, addition of a single DelS31 mutation to FLiRT and KP.2 led to modest yet consistent decreases the titers of FLiRT-DelS31 (p > 0.05) and KP.2_DelS31 (p < 0.01) relative to their parental FLiRT and KP.2 variants, respectively. The exception to this trend was the KP.3.1.1 variant, which contains DelS31 but exhibited a modest increase in titer relative to the parental KP.3 variant (see Discussion) (**Fig 1E**).

### DelS31 decreases nAb titers of LB.1, KP.2.3 and KP.3.1.1 in bivalent mRNA-vaccinated healthcare workers

Next, we sought to elucidate how well these new variants are neutralized by antibodies in several different cohorts of sera (**Fig 2**). First, we investigated a cohort of healthcare workers (HCWs) at the Ohio State University Wexner Medial Center that had received at least two doses of monovalent mRNA vaccine and a dose of the bivalent formulation of the mRNA vaccine that includes both the wildtype and BA.4/5 spikes (n=10) (**Table S1, Fig 2A-B**). Previously, we have shown that JN.1, FLiRT, and KP.2 exhibit modestly reduced nAb titers relative to their ancestral BA.2.86 in this cohort^9^. New JN.1-derived subvariants LB.1, KP.2.3, and KP.3.1.1 all exhibited further decreases in nAb titer of 9.2-fold (p < 0.01), 9.3-fold (p < 0.001), 9.3-fold (p < 0.001) relative to their parental JN.1 variants, respectively. For all variants, this decrease in nAb titer appeared to have been driven by the single DelS31 mutation, which contributed to a decrease of 4.5-fold (p < 0.01), 5.1-fold (p < 0.001), and 5.3-fold (p < 0.001) for FLiRT_DelS31, KP.2_DelS31, and KP.3.1.1 (KP.3 + DelS31) relative to their parental FLiRT, KP.2 and KP.3 subvariants, respectively (**Fig 2A-B**). Once again, single point mutants FLiRT_Q183H and KP.2_H146Q did not have obvious impacts on nAb titer, which showed similar antibody titers to their parental FLiRT and KP.2, respectively (**Fig 2A-B).** Overall, these data show that the DelS31 mutation dictates escape of the newly emerged LB.1, KP.2.3 and KP.3.1.1 variants from bivalent mRNA vaccine-generated nAb responses.

**Figure 2:**
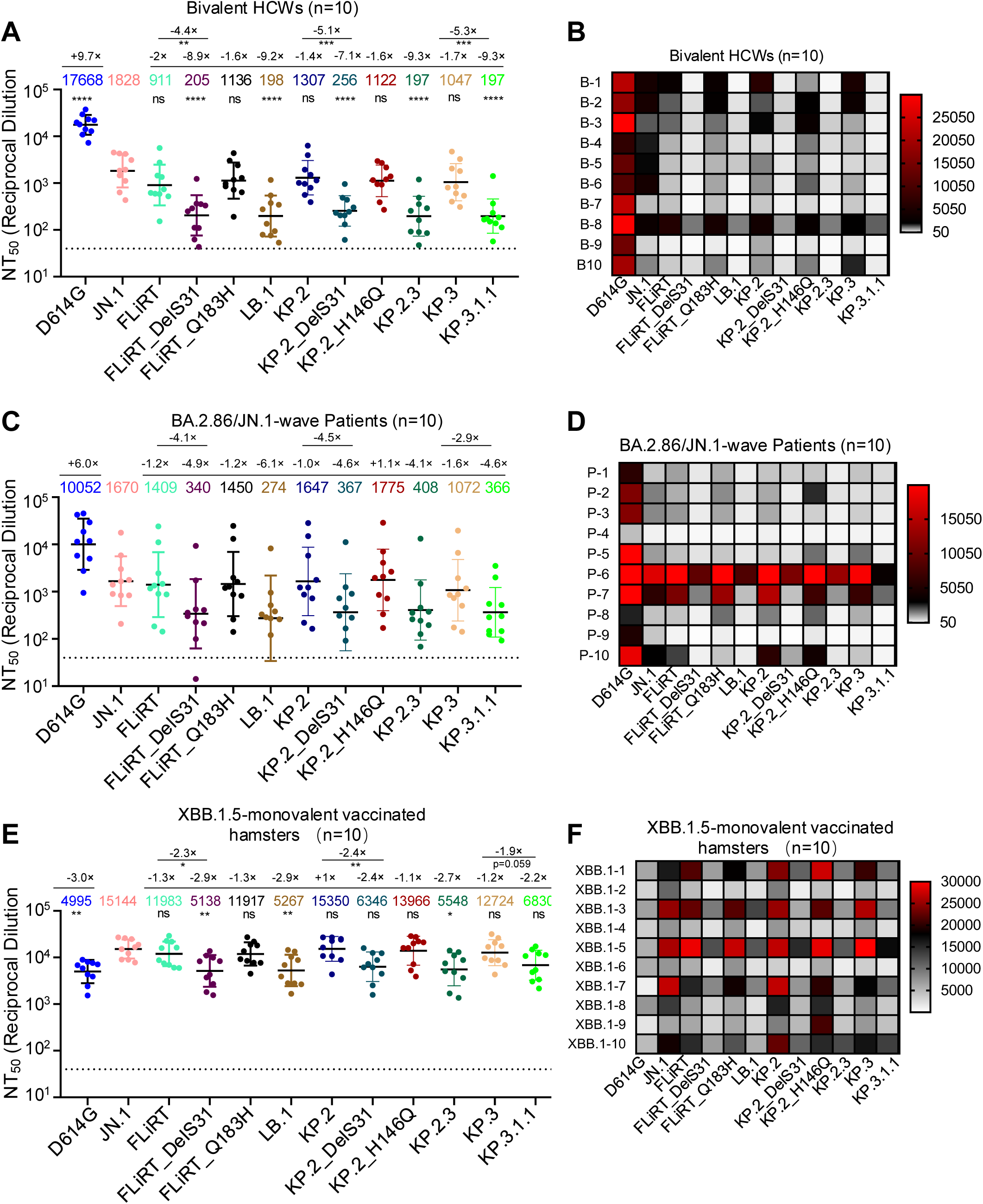
Neutralization of JN.1 variants by antibodies in bivalent-vaccinated HCWs, XBB.1.5-vaccinated hamsters, and BA.2.86/JN.1-infected people. NAb titers were determined against JN.1-derived variants of interest in the sera of **(A-B)** HCWs that received at least two doses of monovalent mRNA vaccine and a dose of bivalent (WT+BA.4/5) mRNA vaccine (n=10), **(C-D)** individuals that were infected during the BA.2.86/JN.1 wave of infection in Columbus, Ohio (n=10), and **(E-F)** golden Syrian hamsters that were vaccinated with two doses of a mumps virus-based monovalent XBB.1.5 spike vaccine (n=10). Plots in **(A, C, and E)** represent geometric mean nAb titers at 50% with standard errors. Geometric mean antibody titers are depicted at the top of the plots with fold changes relative to JN.1 above them. **(B, D and F)** Heatmaps that depict the corresponding nAb values for each cohort listed by individual samples. Significance was determined and displayed relative to JN.1 using log10 transformed values, unless otherwise indicated; stars represent *p < 0.05; **p < 0.01; ****p < 0.0001, and ns p > 0.05.

### DelS31 causes escape of antibodies in BA.2.86/JN.1-wave convalescent sera

We also investigated nAb titers in a cohort of patients admitted to the Ohio State University Wexner Medical Center during the BA.2.86/JN.1 wave of infection in Columbus, Ohio (n=10) (**Table S1**, **Fig 2C-D**). Serum samples from all patients were collected between 1 to 10 days post-infection, and the time from their last vaccination ranged from 34 to 1033 days; all patients had received at least one dose of a monovalent vaccine (**Table S1**). Previously, we have shown that these patients exhibit modestly decreased nAb titers against JN.1-derived subvariants, especially FLiRT, relative to JN.1 likely due to key amino acid mutation R346T and F456L^9^. Here we found that LB.1, KP.2.3, and KP.3.1.1 all exhibited dramatic decreases in nAb titer, with 5.1-fold (p = 0.06), 4.0-fold (p = 0.06), and 2.9-fold (p = 0.09) relative to parental FLiRT, KP.2 and KP.3 variants, respectively. Again, this appeared to be largely driven by the DelS31 mutation, which contributed to decreases in nAb titer of 4.1-fold (p < 0.07), 4.5-fold (p = 0.07), and 2.9-fold (p = 0.09) for FLiRT_del31, KP.2_del31, and KP.3.1.1 relative to parental FLiRT, KP.2 and KP.3 variants, respectively (**Fig 2C-D**). Note that the greater p values presented here compared to those shown above for bivalent serum samples was likely due to large variations among patients of these two cohorts—one being first responders who became COVID positive and suffered mild illness (n = 4), and another being ICU patients with large differences in age and clinical conditions (n = 6) (**Table S1**, **Fig. S2A-B**). In particular, we noted that patients P6 and P7 in the ICU group exhibited higher titers against the JN.1-lineage variants, especially DelS31. P6 was a 77-year-old man, who received a single dose of the Moderna monovalent vaccine followed by one dose of the Pfizer bivalent vaccine; his samples were collected 434 days after his final vaccination. P7, on the other hand, was a 46-year-old woman, who was administered three doses of the Moderna monovalent vaccine and one dose of the Moderna bivalent vaccine; her samples were taken 334 days after her last vaccination. P10 also showed relatively high titers for D614G, though with similar trends of decrease for JN.1 and JN.1-derived subvariants (**Fig 2C-D**). Collectively, these results are in accordance with the patten of bivalent mRNA-vaccinated sera, supporting the conclusion that DelS31 drives antibody escape.

### XBB.1.5 monovalent-vaccinated hamster sera robustly neutralize JN.1 variants, with reduced titers for subvariants harboring DelS31 in the spike

The last cohort of sera we investigated was golden Syrian hamsters that were administered two doses of a recombinant mumps virus-based monovalent XBB.1.5 spike vaccine (**Fig 2E-F**). The nAb titer in this group was the highest against JN.1, with modest reductions for FLiRT and KP.2 as we have shown previously^9^. Even with this strong response, we observed clear decreases in nAb titers for LB.1, KP.2.3, and KP.3.1.1, with 2.3-fold (p < 0.05), 2.8-fold (p < 0.01), and 1.9-fold (p = 0.059) decreases relative to parental FLiRT, KP.2, and KP.3 variants, respectively. Similar to other cohorts described above, this decrease was driven by the DelS31 mutation, with FLiRT_DelS31, KP.2_DelS31, and KP.3.1.1 exhibiting decreases of 2.3-fold (p < 0.05), 2.4-fold (p < 0.01), and 1.9-fold (p = 0.059) relative to parental FLiRT, KP.2 and KP.3 variants, respectively (**Fig 2E-F**). Altogether, results from these three cohorts reveal an essential role for DelS31 located in the NTD of spike in driving nAb escape from vaccination and infection.

### Monoclonal antibody S309 is completely ineffective against JN.1-derived variants

We have previously shown that BA.2.86-derived Omicron variants, including JN.1 and subvariants, are resistant to neutralization by class 3 monoclonal antibody S309^8,9,14^, one of the most broadly neutralizing antibodies characterized^22–25^. Here we found that all newly emerged JN.1 subvariants, including LB.1, KP.2.3 and KP.3.1.1, also were completely resistant to neutralization by S309, with calculated IC_50_ values similar to their ancestral FLiRT, KP.2 and KP.3 variants (**Fig 3A-B**).

**Figure 3:**
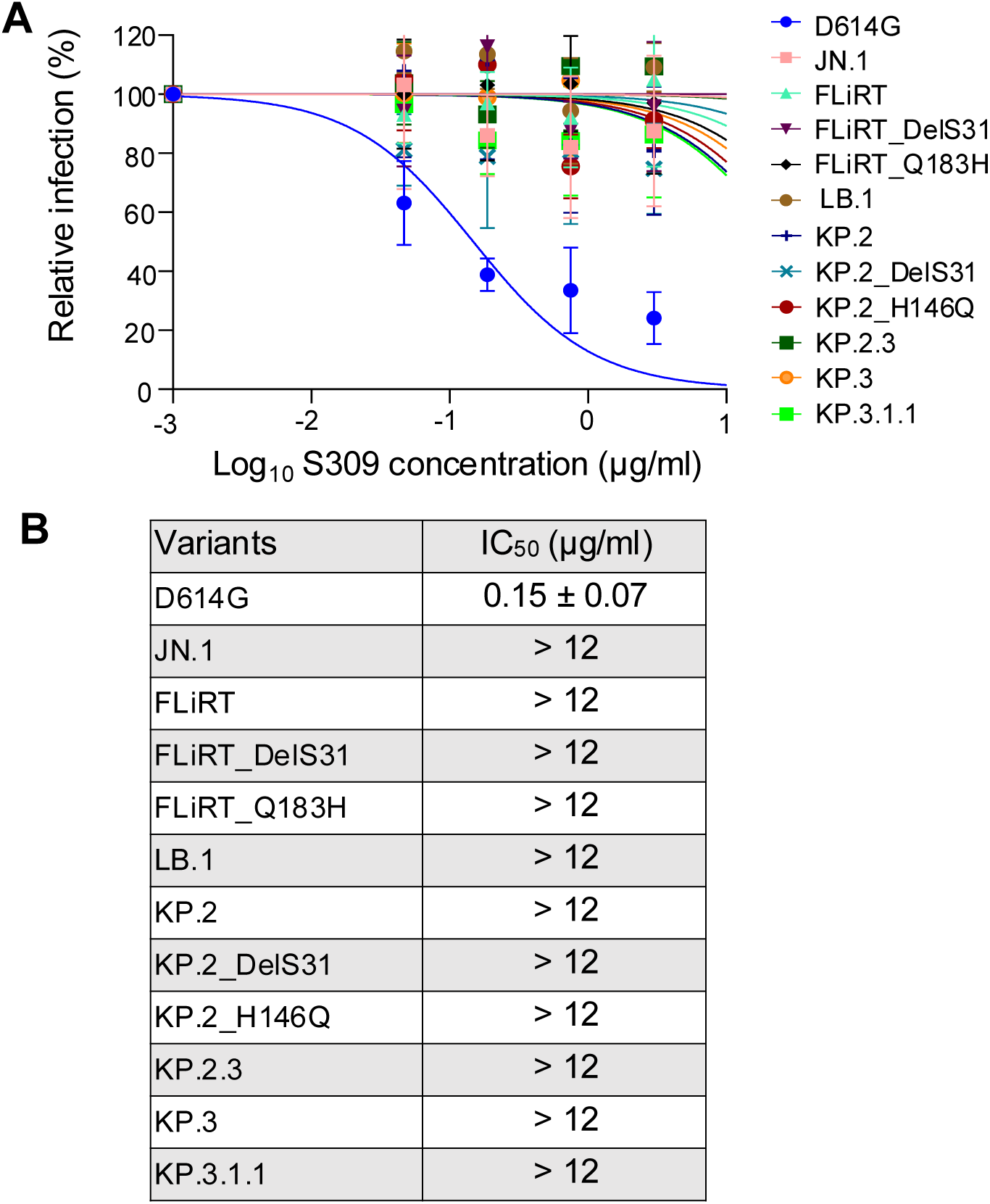
Neutralization of JN.1 variants by monoclonal antibody S309. Neutralization by class 3 monoclonal antibody S309 was determined for JN.1-derived variants of interest and plotted **(A),** and inhibitory concentrations at 50% (IC_50_) was determined and displayed in **(B)**. Raw luminescence values were normalized to untreated controls for plotting and IC_50_ calculations.

### DelS31 drives antigenic differences of newly emerged JN.1 subvariants from their parental variants

To further supplement our nAb data, we conducted antigenic mapping analyses to determine the relative antigenic distances between each variant for the three cohorts of sera. Overall, trends of antigenic distances between the three cohorts were comparable, with XBB.1.5-vaccinated hamster sera having much smaller antigenic distances between variants than the bivalent-vaccinated people and BA.2.86/JN.1-wave patients as we have demonstrated previously^9,14^ (**Fig. 4A-C**). Importantly, all subvariants possessing the DelS31 mutation, i.e., FLiRT_DelS31, LB.1, KP.2_DelS31, KP.2.3, and KP.3.1.1 clustered together and were distinct from the other JN.1-derived subvariants (**Fig 4A-C**) – with relatively longer antigenic distances to JN.1 (4.7 ∼ 5.6 AU) compared to their ancestral FLiRT, KP.2 and KP.3 (1.0 ∼ 1.9 AU) (**Fig 4D**, **Fig. S2A-C**). Overall, the antigenic data are consistent with the patterns of neutralization for each variant, highlighting once again the crucial role of a single DelS31 mutation, which dictates the antibody escape and shapes the antigenicity of the spike protein.

**Figure 4:**
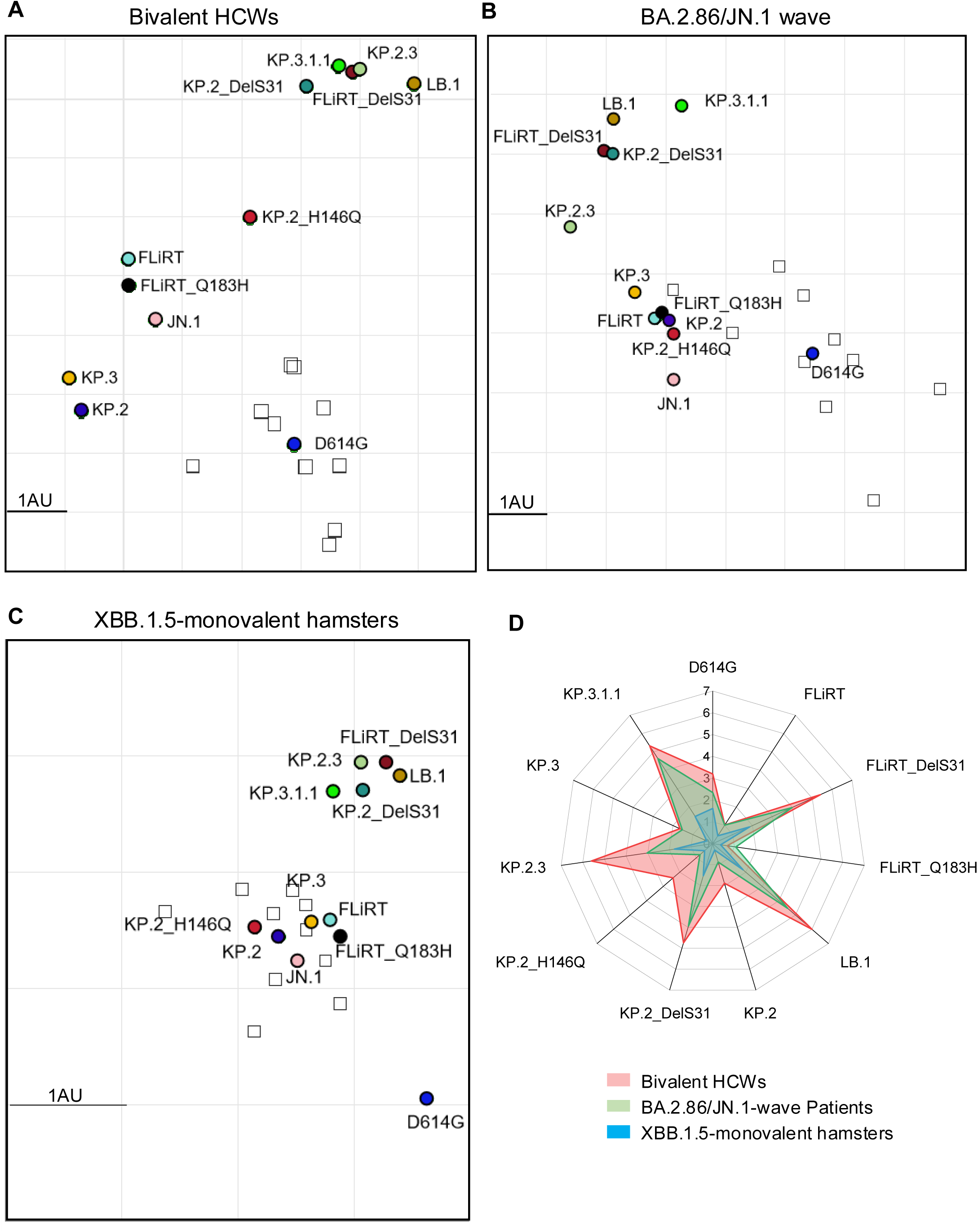
Antigenic mapping of neutralization data against JN.1 variants. The Racmacs program was used to plot relative antigenic distances between each spike antigen (circles) and sera sample (squares) for **(A)** the bivalent-vaccinated HCWs, **(B)** the BA.2.86/JN.1-wave infected people, and **(C)** the XBB.1.5-vaccinated hamsters. The scale bar represents 1 antigenic distance unit (AU) which is equivalent to about a 2-fold different in nAb titer. **(D)** The antigenic distances of each variant relative to JN.1 from three groups of cohorts (n=3) were averaged and plotted. The scale bar represents 1 antigenic distance unit (AU).

### DelS31 decreases cell-cell fusion mediated by the spike

We sought to better understand how N-terminal mutations impact different aspects of spike protein biology. First, we investigated the spike’s ability to trigger fusion between cell membranes in a syncytia formation assay wherein 293T cells transfected with spike were co-cultured with target 293T-ACE2 cells (**Fig 5A-B**) or CaLu-3 cells (**Fig 5C-D**). Consistent with the pattern of all Omicron variants^4,8,9,14,20,26,27^,and similar to their ancestral FLiRT, KP.2 and KP.3, all newly emerged JN.1 subvariants exhibited decreased fusion relative to D614G and JN.1 in both cell lines (**Fig. 5A-D**). Notably, we found that, although fusion mediated by LB.1, KP.2.3, and KP.3.1.1 was comparable to the parental variant JN.1, DelS31 variants harboring the single DelS31 mutation consistently showed decreased levels of cell-cell fusion compared to their parental FLiRT, KP.2 and KP.3 variants in 293T-ACE2 cells. Specifically, FLiRT_DelS31, KP.2_DelS31, and KP.3.1.1 (i.e, KP.3_DelS31) variants exhibited decreases of 1.2-fold (p < 0.01), 1.3-fold (p < 0.01), and 1.2-fold (p < 0.001) relative to their parental FLiRT, KP.2 and KP.3 variants, respectively (**Fig 5A-B**). In contrast, a single Q183H mutation in FLiRT or H146Q in KP.2 did not have any impact on cell-cell fusion of LB.1 and KP.2.3 (**Fig 5A-B)**. Similar results were obtained in CaLu-3 cells, where decreases of 1.3-fold (p < 0.01), 1.1-fold (p < 0.01), and 1.2-fold (p < 0.001) were found for FLiRT_DelS31, KP.2_DelS31, and KP.3.1.1 relative to their parental FLiRT, KP.2 and KP.3 variants, respectively (**Fig 5C-D**). These results suggest that the DelS31 mutation at the NTD significantly decreases fusion mediated by the SARS-CoV-2 spike, at least in JN.1-derived Omicron subvariants.

**Figure 5:**
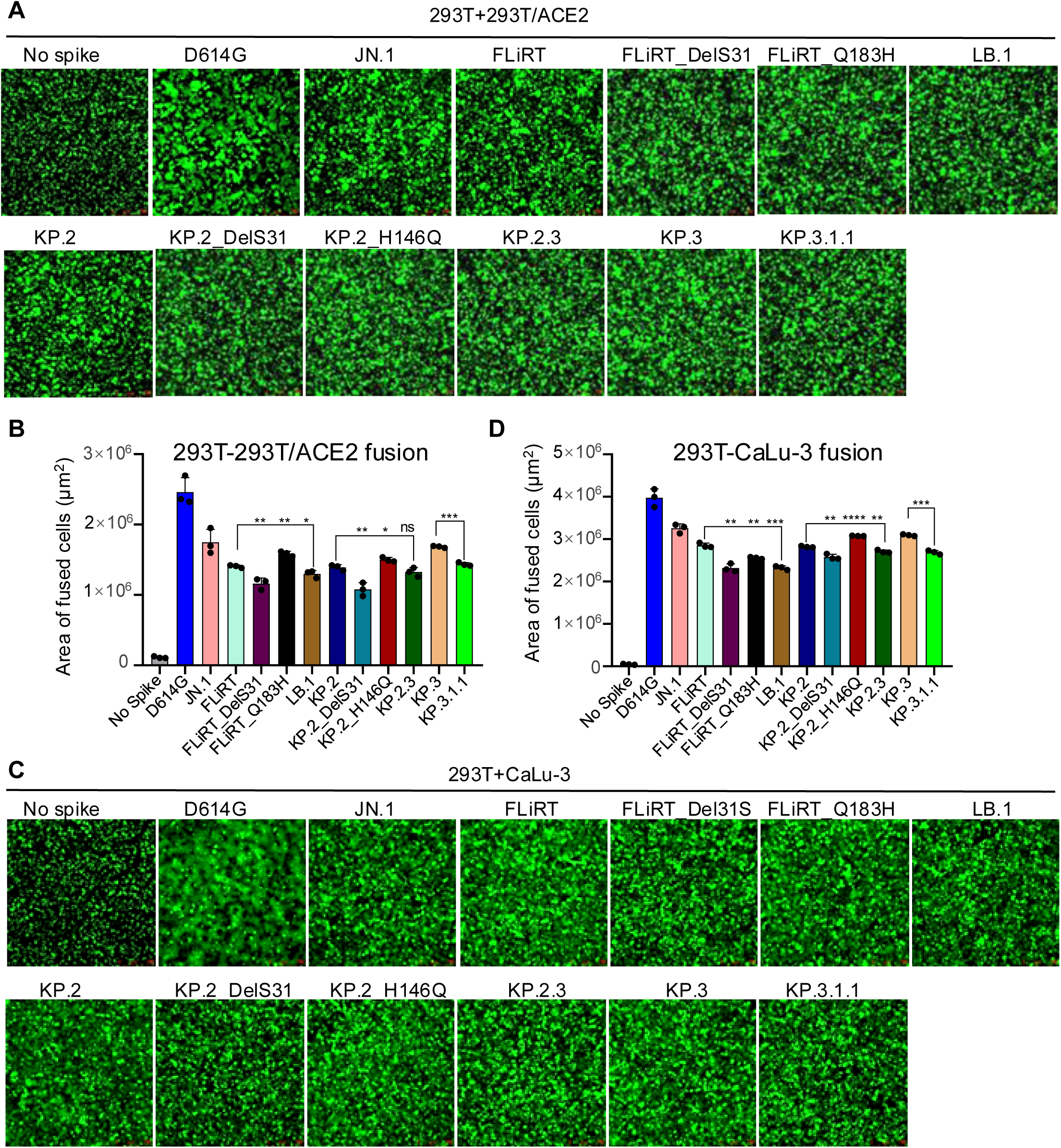
Cell-cell fusion of JN.1-derived spikes. Fusion triggered between membranes by the spike proteins of interest was determined between 293T cells expressing the spike and 293T-ACE2 cells overexpressing ACE2 or CaLu-3 cells expressing an endogenous level of ACE2. Representative images of fusion are depicted for **(A)** 293T-ACE2 and **(C)** CaLu-3 and quantification of total areas of fusion across 3 images are represented for **(B)** 293T-ACE2 and **(D)** CaLu-3. Bars represent means with standard deviation, significance was determined relative to ancestral variants as indicated, and stars represent *p < 0.05; **p < 0.01; ***p < 0.001; ****p < 0.0001 and ns: p > 0.05.

### DelS31 variants exhibit increased surface expression despite comparable processing

Another critical aspect of spike biology is its ability to be expressed on the plasma membrane following intracellular cleavage and trafficking. Importantly, this feature is directly associated with membrane fusion activity. We assessed this feature by performing surface staining against the S1 subunit of spike on 293T cells producing pseudotyped lentiviruses. We found that all JN.1-derived variants, including newly emerged KP.3, LB.1, KP.2.3 and KP.3.1.1 subvariants, had decreased levels of surface expression relative to D614G, similar to what we have shown for JN.1 and prior JN.1 variants such as FLiRT and KP.2^9,14^. Notably, variants that harbor the single DelS31 mutation, i.e., FLiRT_DelD31, KP.2_DelS31, and KP.3.1.1, showed 20∼30% increased expression on the cell surface relative to their parental FLiRT (p < 0.0001), KP.2 (p < 0.001) and KP.3 (p < 0.0001), respectively— based on their calculated geometric means (**Fig 6A-B**). In contrast, the single mutations Q183H and H146Q, which are also located in the NTD of spike, did not affect the cell surface expression of LB.1 and KP.2.3 subvariants (**Fig 6A-B)**.

**Figure 6:**
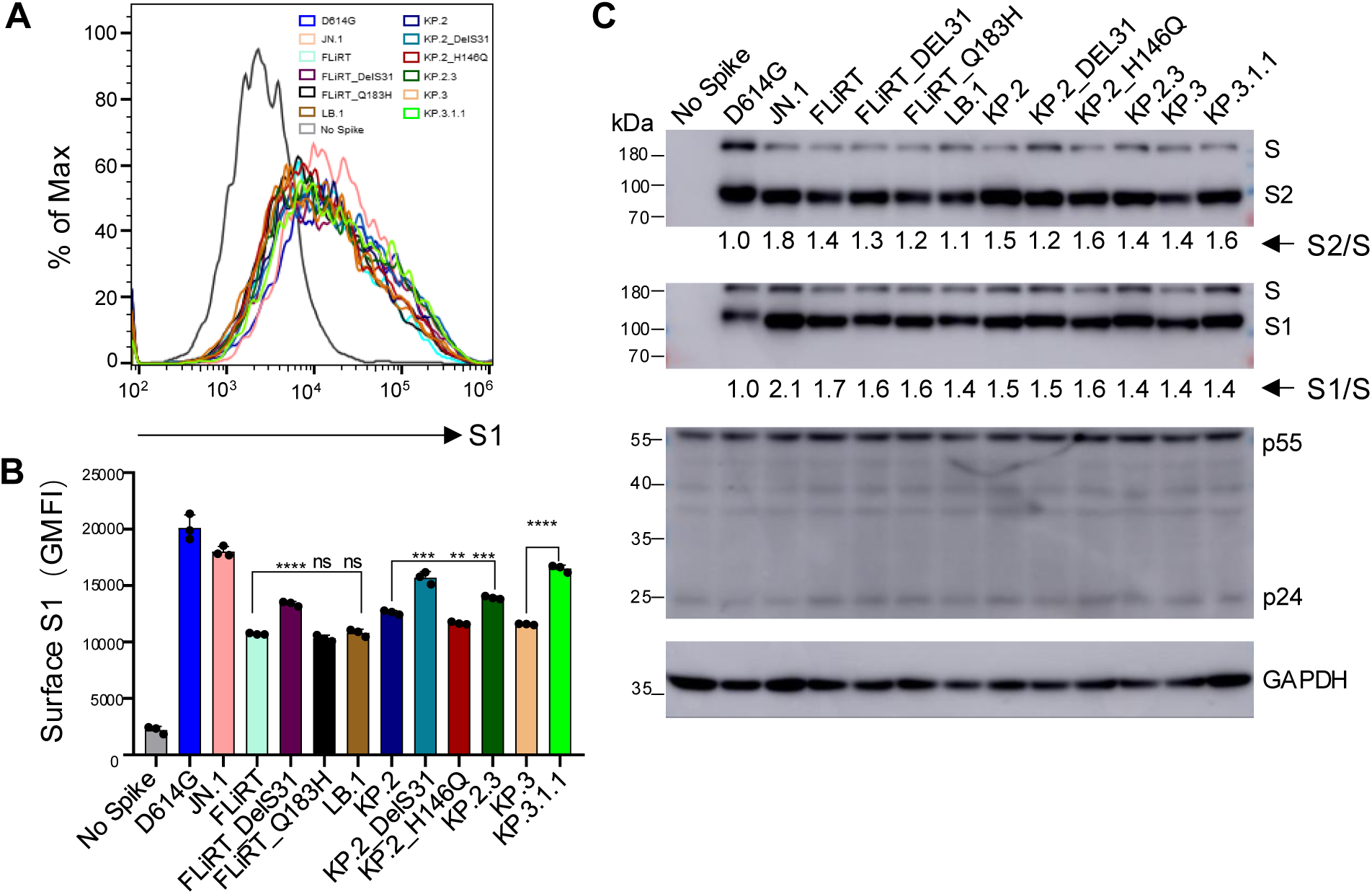
Surface expression and processing of JN.1-derived spikes. **(A-B)** The surface of 293T cells used to produce pseudotyped vectors was probed with anti-S1 antibody to compare surface expression between spikes of interest. **(A)** Representative histograms depicting surface expression and **(B)** geometric mean intensities (MFIs) of surface S1 are depicted (n=3). **(C)** Processing of spikes into S1/S2 subunits by furin was determined by lysing 293T cells used to produce pseudotyped viruses and probed by using anti-S1, anti-S2, anti-p24, and anti-GAPDH antibodies. Relative ratios of S2/S or S1/S were quantified using NIH ImageJ, calculated by comparing to D614G, and are displayed under corresponding blots. The plot in **(B)** represents geometric means with standard deviation and significance was determined relative to parental FLiRT, KP.2 or KP.3 variants as indicated; stars represent **p < 0.01; ***p < 0.001; ****p < 0.0001, and ns: n > 0.05.

We next performed western blotting to determine the ability of the spike protein to be processed by furin in virus-producer cell lysates by quantifying the ratios of S1 and S2 subunits vs. full-length spikes. When compared to D614G, JN.1 exhibited a marked increase in processing, whereas FLiRT and KP.2 showed a decrease relative to JN.1, as we have shown previously^9^ (**Fig 6C**). Of note, the new JN.1 subvariants LB.1, KP.2.3 and KP.3.1.1, alongside their DelS31 and other single mutants, did not exhibit obvious changes in spike processing, remaining similar to their parental FLiRT, KP.2 and KP.3 variants, respectively (**Fig 6C**). Comparable transfection efficacy and cell lysis was confirmed by similar signals of HIV-1 Gag and cellular GAPDH (**Fig 6C**) Interestingly, all DelS31-containing S1 signals migrated slower than other JN.1-derived variants, likely due to the acquisition of a potential N-linked glycosylation site (see Discussion).

### DelS31 stabilizes spike of newly emerged JN.1 subvariants

The above results from viral infectivity, membrane fusion, and cell surface expression assays suggest that the DelS31 mutation may confer increased conformational stability to the spike protein of newly emerged JN.1 subvariants compared to their ancestral forms. To test this directly, we incubated purified pseudotyped viruses, with or without the DelS31 mutation, at temperatures of 37°C, 39°C, 41°C and 43°C for 1 hour and assessed their impact on viral infectivity by infecting 293T/ACE2 cells. Viruses kept at 4°C for the same period of time served as a control. As expected, incubation at elevated temperatures (37 to 43°C) gradually reduced the infectivity of lentiviral particles bearing the spike proteins of these variants (**Fig 7A-B**). Notably, JN.1 subvariants with the DelS31 mutation, particularly KP.3.1.1 (KP3_DelS31) and FLiRT_DelS31, demonstrated greatly increased resistance to temperature-induced inactivation, with calculated half-lives (T1/2) of 41.68°C (± 0.14) and 39.25°C (± 0.04), respectively. This was in contrast to their parental KP.3 and FLiRT variants, which had T_1/2_ values of 39.12°C (± 0.02) and 37.23°C (± 0.08) (**Fig 7A-B**). Among all variants examined, FLiRT exhibited the least stability, i.e., T_1/2_ of 37.23°C (± 0.08), while KP.3-DelS31 showed the greatest stability (**Fig 7A-B**).

**Figure 7:**
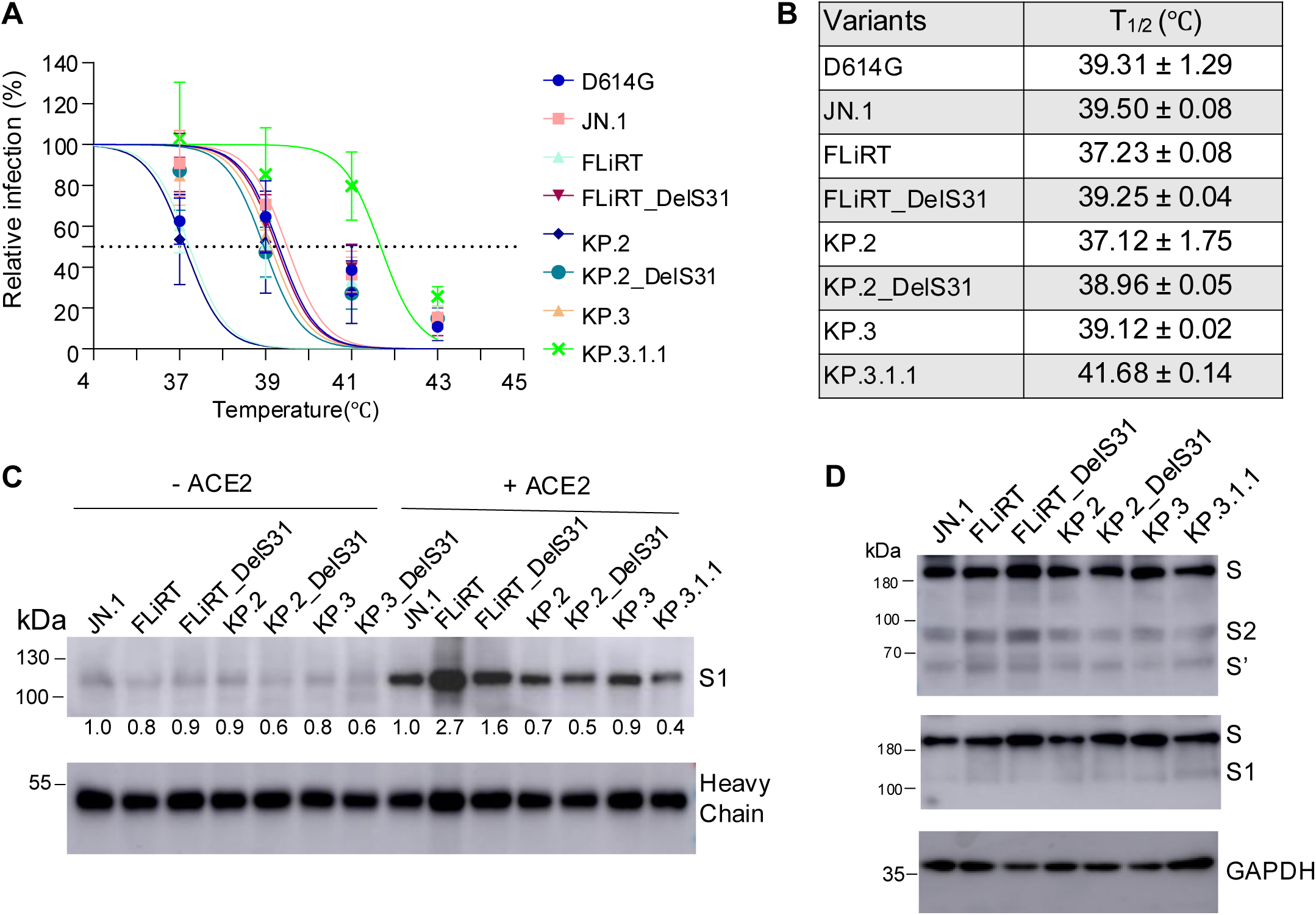
Stability of JN.1 variant spikes and pseudotyped viral particles. (**A**) Lentiviral pseudovirions were purified (without serum) and incubated at indicated temperatures (37 to 43°C) for 1 h, and viral infectivity was determined by infecting 293T-ACE2 cells. Relative percent of infection is plotted by comparing the titer at 4°C, which was set to 100%. For each variant, the temperature at which the viral infectivity was lost by 50% (T_1/2_) was determined and displayed by a dashed line in **(A)** and listed in **(B)**. (**C**) HEK293T cells were transfected with spike constructs of interest and treated with or without sACE2 (10 µg/ml) for 4 h. Cell culture media and lysates were collected, with shed S1 proteins being immunoprecipitated with an anti-S1 antibody. **(D)** Cell lysates were blotted with anti-S2, anti-S1 and anti-GAPDH antibodies, and relative signals were quantified by NIH ImageJ by setting the value of JN.1 to 1.0.

Next, we assessed S1 shedding of these JN.1 subvariants by transfecting 293T cells with the spike protein of interest, followed by treatment with or without 10 µg/ml of soluble ACE2 (sACE2). The culture media and cell lysates were harvested and immunoblotted with an anti-S1 antibody. As shown in **Fig 7C**, sACE2 treatment significantly stimulated S1 shedding of all spikes examined, validating the experimental procedure. Interestingly, variants containing the DelS31 mutation, especially KP.2-DelS31 and KP.3.1.1 (KP.3_DelS31), exhibited decreased levels of S1 shedding compared to their parental KP.2 and KP.3 variants, both in the presence and absence of sACE2 (**Fig 7C**). In the case of FLiRT-DelS31 and FLiRT, while the former showed reduced shedding compared to the latter, no significant difference was observed between them in the absence of sACE2 (**Fig 7C**). The antibody heavy chain signals were consistent across samples, indicating equal amounts of anti-S1 antibody were used to pull down the S1 protein from the culture media. Lysates of transfected and sACE2-treated cells were blotted with anti-S1 and anti-S2 antibodies, showing comparable levels of spike expression and cleavage into S1 and S2 (**Fig 7D**). In the cell lysates, we observed significantly reduced S1 signals and enhanced S2’ intensity, the latter being an indicator of spike activation (**Fig 7D**). This was consistent with the transfected cells being activated by sACE2 before lysis, leading to increased S1 shedding from the cell surface as well as enhanced S2 cleavage upon sACE2 engagement.

### Molecular modeling of key NTD mutations in LB.1, KP.2.3, and KP.3.1.1 spikes

To better understand the underlying mechanisms, especially the impact of spike mutations on these new variants, we performed homology modeling to investigate alterations in receptor engagement, spike conformational stability, and antibody interactions. The DelS31 mutation causes a positional shift in the adjacent residue F32, orienting it towards the core of the NTD and enabling it to form strong hydrophobic interactions with surrounding core residues, including T29, R34, V62, L56, Y91, and F216 (**Fig 8A**). Compared to the original unfavorable polar-to-hydrophobic interaction mediated by S31, this serine-to-phenylalanine substitution enhances the stability of the NTD and induces a conformational change that reshapes the domain. This alteration likely strengthens the interaction between the NTD and the RBD, leading the RBD to energetically favor the down conformation (**Fig 8B**). As a result, the receptor-binding motif (RBM) becomes less accessible to both ACE2 receptor binding (**Fig 8C**) and some neutralizing antibodies, such as RBM-targeting class 1 and inner face-targeting class 4 antibodies. On the contrary, recognition of class 2 and 3 antibodies is not affected by this mechanism (**Fig 8D**). The other mutations observed in KP.2.3, and LB.1 spike, such as H148Q and Q183H, disrupt the epitope region of some NTD-targeting neutralizing antibodies, including 4A8 **(Fig 8E**) and C1520 (**Fig 8F**). Mutation of these residues presumably disrupts the binding of these antibodies to spike. Additionally, the DelS31 mutation introduces an N-linked glycosylation sequon (<u>N</u>F<u>T</u>), resulting in a glycan modification at the N30 residue. Together with the adjacent N-linked glycan at N61, these glycan chains may interact with each other, effectively shielding the underlying region of the NTD from antibody recognition (**Fig 8A**, **Fig 8G)**.

**Figure 8:**
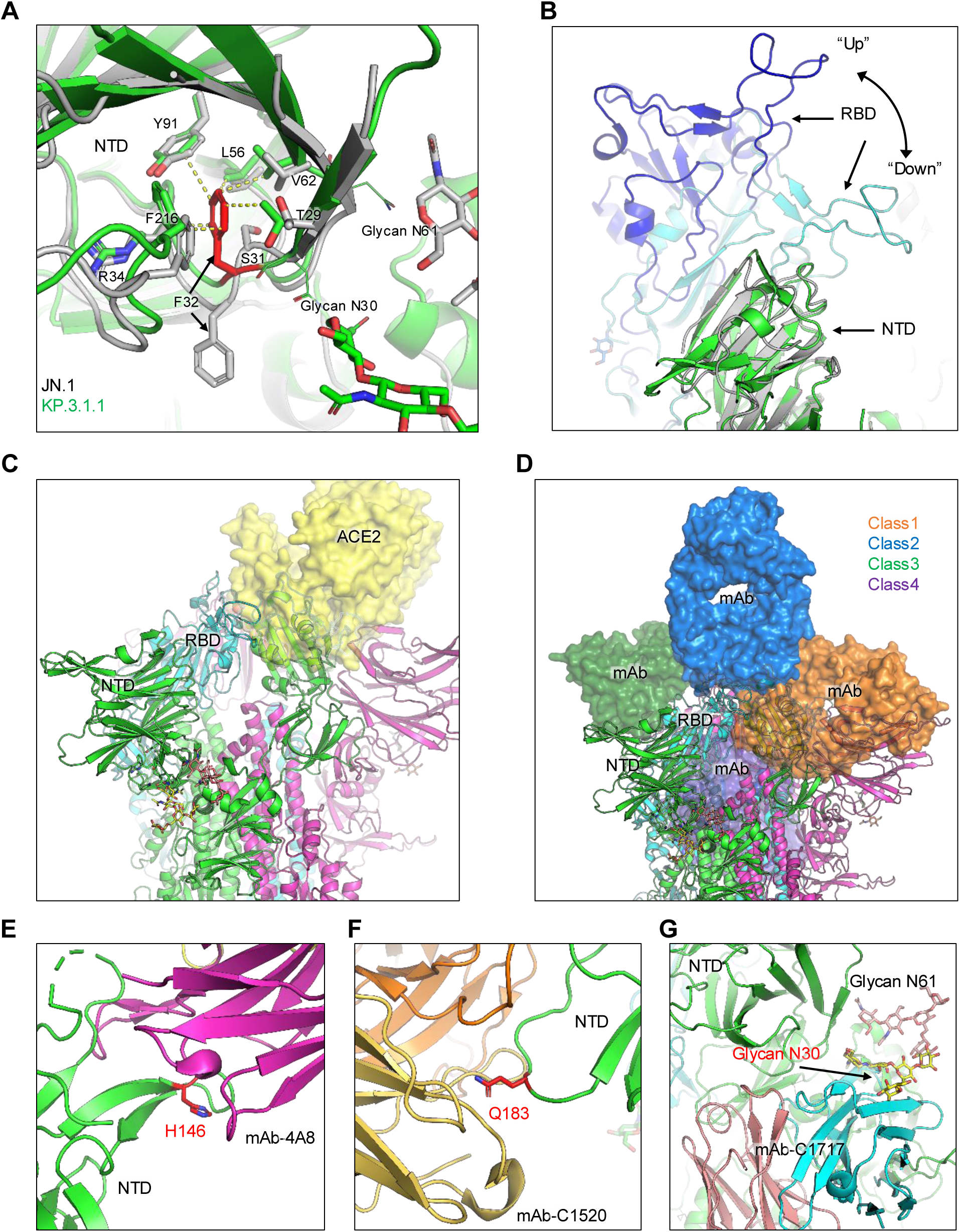
Structural modeling of key NTD mutations in LB.1, KP.2.3, and KP.3.1.1. **(A)** Structural comparisons of NTD between KP.3.1.1 and JN.1 spike proteins. The DelS31 mutation shifts F32, thus altering its side chain direction to form hydrophobic interactions with surrounding NTD core residues, including T29, R34, V62, L56, Y91, and F216, while introducing glycosylation at N30. **(B)** The DelS31 mutation stabilizes the NTD, reshaping its conformation and enhancing its interaction with the receptor-binding domain (RBD) to favor the RBD down conformation. **(C)** The down conformation reduces RBD accessibility to the ACE2 receptor (yellow surface). **(D)** This down conformation restricts the accessibility of class 1 and 4 antibodies, but not class 2 and 3 antibodies. Antibodies are depicted as semi-transparent surfaces. **(E)** and **(F)** Mutations at residues (shown as sticks) H146 and Q183 disrupt the epitopes of certain NTD-targeting antibodies, such as 4A8 and C1520. **(G)** Glycosylation (shown as sticks) at N30 interferes with the recognition of some NTD-targeting antibodies, such as C1717.

## DISCUSSION

The ongoing evolution of SARS-CoV-2 presents a significant challenge to the sustained control of the COVID-19 pandemic. The emergence of the JN.1 variant raised new concerns due to its pronounced immune evasion and higher transmissibility compared to its ancestral variant, BA.2.86. The convergent evolution of key residues in the spike protein of early JN.1-derived subvariants—such as R346, L455, and F456 in the RBD—has further exacerbated immune escape, underscoring an urgent need for updated vaccine formulations as cases surged during the summer of 2024^29^. As summer draws to a close, several new subvariants from the JN.1 lineage are competing for dominance globally. Among these, the LB.1, KP.2.3, and KP.3.1.1 variants, which collectively account for over 90% of COVID-19 cases, are characterized by mutations concentrated in the NTD of the spike protein (**Fig. 1**). Notably, these variants have convergently acquired the DelS31, suggesting that this mutation may confer a fitness advantage.

Our study demonstrates that this key mutation significantly contributes to spike protein and viral particle stability, enhanced evasion of neutralizing antibodies, distinct antigenicity, and reduced cell-cell fusion. The observation of such distinct phenotypes resulting from a single NTD mutation, particularly in neutralization and spike stability, is striking. Notably, the reduction in neutralization conferred by DelS31 in convalescent sera aligns with findings from other studies^30–32^, though the decreases observed in our cohorts were more pronounced. This difference may be attributed to the fact that most cohorts in other studies had been repeatedly exposed to Omicron variants, which likely boosted antibody responses against more recently circulating variants and helped mitigate immune imprinting from early pandemic strain^22,33,34^. Importantly, we found that variants harboring the DelS31 mutation are antigenically distinct from other JN.1 subvariants across all cohorts analyzed (**Fig 4**). Together, our results highlight the necessity of developing new vaccine antigens that incorporate the DelS31 mutation to effectively curb the pandemic, in addition to the existing JN.1 and KP.2 formulations recently approved by the US FDA.

Our study revealed that newly emerged JN.1 subvariants, including LB.1, KP.2.3, and KP.3.1.1, are completely resistant to neutralization by S309, one of the most broadly neutralizing antibodies tested^22–24,35,36^. While this finding aligns with expectations—since these subvariants retain the conserved S309-binding motifs from their ancestral BA.2.86 and JN.1 variants^8,14,20,37–39^—it underscores the urgent need for the development of new antibodies capable of effectively neutralizing these evolving variants to maintain control over the COVID-19 pandemic^40^. Recent studies on newly approved monoclonal antibodies Pemivibart, Sipavibart, and SA55 have shown that while these antibodies were initially effective as prophylactic measures against some recent variants^41–43^, Pemivibart and Sipavibart have lost activity against KP.2, LB.1, and KP.3.1.1^44,45^. Fortunately, SA55 has retained its efficacy against most of these newly emerged JN.1 subvariants^43^. Overall, our findings suggest that the DelS31 mutation plays a significant role in reducing the efficacy of these monoclonal antibodies, despite being distant from their epitopes, indicating possible epistatic effects on neutralization (**Fig 8**). They also highlight the critical need for ongoing development of novel antiviral drugs for both prevention and therapy.

The spike residue S31 has not been extensively studied due to the absence of mutations in prior variants. However, research by the Bloom lab using high-throughput screens of spike mutations that could impact ACE2 affinity or antibody neutralization identified S31 as a residue of potential interest^17^. Their study demonstrated that mutations at S31 could reduce ACE2 binding in the context of the XBB.1.5 variant, likely due to conformational changes that favor the spike occupying an RBD-down conformation more frequently. Our molecular modeling reveals that the DelS31 mutation enhances the stability of the NTD by reorienting the adjacent phenylalanine residue from an outward to an inward position (**Fig 8**). This reorientation creates energetically favorable hydrophobic interactions with the NTD core. As a result, the NTD is stabilized and undergoes a slight conformational change that strengthens its interaction with the RBD, favoring the “down” conformation of the RBD. This “down” conformation reduces the accessibility of the receptor-binding motif (RBM) to both the ACE2 receptor and certain neutralizing antibodies. Additionally, mutations in KP.2.3 and LB.1 disrupt epitopes targeted by some NTD-specific antibodies. The DelS31 mutation also introduces glycosylation at N30, which helps shield the NTD from antibody recognition. Detailed structural studies of DelS31 will be necessary to confirm these potential changes in spike conformation, particularly concerning ACE2 engagement and antibody recognition.

Of particular interest is the impact of the DelS31 mutation on infectivity, cell-cell fusion, and cell surface expression. We observed that DelS31 increased infectivity in 293T-ACE2 cells, a finding corroborated by another study that reported similar results for KP.3.1.1 compared to KP.3 in HOS-ACE2-TMPRSS2 cells^30^. However, in CaLu-3 cells, the DelS31 mutation led to reduced viral infectivity and impaired cell-cell fusion (**Fig 1**, **Fig 5**). We speculate that this difference is due to the lower levels of ACE2 expression in CaLu-3 cells, as opposed to the overexpression of ACE2 in 293T-ACE2 cells (see modeling in **Fig 8**). Noticeably, fusion is reduced despite the relatively high level of surface expression of DelS31 variants (**Fig. 5**, **Fig. 6**). The reduced fusion is likely driven by the fact that the spike cannot as readily engage with ACE2, evidenced by the decreased S1 shedding of DelS31 variants compared to their ancestral variants (**Fig 7**). Our molecular modeling analysis suggests that DelS31 stabilizes the spike protein by promoting an RBD-down conformation, which could help prevent premature triggering for membrane fusion^46,47^. Interestingly, KP.3.1.1 shows slightly higher infectivity than KP.3 in CaLu-3 cells (**Fig. 1E**), and this could be attributed to the presence of the Q493E mutation, which co-occurs with DelS31 in KP.3.1.1. Historically, the Q493E mutation negatively impacted infectivity in previous variants^31,48,49^. However, recent studies suggest that when Q493E co-occurs with L455S and F456L, it enhances ACE2 binding, likely compensating for the reduced binding caused by DelS31^31,49^. This reflects the virus’s ongoing evolutionary trade-offs between ACE2 binding/infection and immune escape.

In addition to our modeling work, we provide experimental evidence that the spike protein of DelS31 variants is more conformationally stable (**Fig 8**). This stability is demonstrated by decreased S1 shedding into culture media—both with and without sACE2 stimulation—as well as increased resistance to elevated temperatures (**Fig 7**). While these findings align with the observed decreased infectivity and impaired cell-cell fusion of these DelS31 variants in CaLu-3 cells (**Fig 1**, **Fig 5**), they could also potentially explain, in part, their dominance during the summer of 2024, in particular KP.3.1.1. More importantly, these findings underscore the role of spike protein stability as a key factor driving viral evolution and fitness, and is also a reminiscent of the original Omicron BA.1 variant, which is more stable compared to its ancestral D614G, Delta, and some earlier variants^50^. It should be noted that while the use of spike and pseudotyped viruses offers clear and direct evidence of the spike’s role, further studies comparing DelS31 variants with those lacking this mutation in the context of authentic viruses will provide additional insights into the mechanisms at play.

Overall, our findings highlight the significant changes in spike biology that can result from a single mutation, particularly one located in the NTD. Our results, along with those from other studies^9,30–32,44^, support the U.S. FDA’s decision to select JN.1/KP.2 as the spike for the latest mRNA vaccine formulation^51^. However, our data also suggest that vaccines incorporating DelS31-containing spikes should be considered as potential immunogens. Additionally, our study underscores the importance of ongoing surveillance of circulating variants to inform pandemic control measures, including vaccination strategies.

## LIMITATIONS OF STUDY

Our study makes use of pseudotyped viruses bearing the SARS-CoV-2 spikes of interest, or the spike protein alone in transfected cells, but lacks analyses with live authentic SARS-CoV-2. We have previously validated our pseudotyped infectivity and neutralization assays alongside authentic SARS-CoV-2^52^ and believe that the timeliness of this data justifies the use of lentiviral pseudotypes. The application of spikes in transfected cells simplifies the study system and allows us to pinpoint the unique role of spike in nAb evasion and conformational stability. We also want to note that our cohorts used in the neutralization assays are relatively small. However, we have previously published using similarly sized cohorts^9,14^ and similar cohort sizes have been used by corroborating studies^30,31^, so we therefore believe these results contribute meaningfully to the discussion of antibody evasion by these variants.

## ACKNOWLEDGEMENTS

We thank the Clinical Research Center and Center for Clinical Research Management of The Ohio State University Wexner Medical Center and The Ohio State University College of Medicine in Columbus, Ohio, especially Breona Edwards, Evan Long, J. Brandon Massengill, Francesca Madiai, Dina McGowan, and Trina Wemlinger, for collecting and processing the samples. We also thank Tongqing Zhou at NIH’s Vaccine Research Center for providing the S309 monoclonal antibody. In addition, we thank Sarah Karow, Gabrielle Swoope, Rushil Madan, and Kristina Luikart of the Critical Care Clinical Trials team of The Ohio State University for sample collection and other supports. We appreciate the assistance by Moemen Eltobgy in sample processing. We also acknowledge Ashish R. Panchal, Mirela Anghelina, Soledad Fernandez, and Patrick Stevens for their assistance in providing the sample information of the first responders and their household contacts. We thank Peng Ru and Lauren Masters for sequencing and Xiaokang Pan for bioinformatic analysis. S.-L.L., D. J., R.J.G., L.J.S. and E.M.O. were supported by the National Cancer Institute of the NIH under award no. U54CA260582. The content is solely the responsibility of the authors and does not necessarily represent the official views of the National Institutes of Health. This work was also supported by a fund provided by an anonymous private donor to OSU. K.X. was supported by NIH grants U01 AI173348 and UH2 AI171611. J.N.F. was supported by a Glenn Barber Fellowship from the Ohio State University College of Veterinary Medicine. M.C. was supported by an NIH T32 training grant (T32AI165391) and Dean’s Graduate Enrichment Fellowship at The Ohio State University. J. L. was supported by NIH R01AI090060 and P01AI175399. J.S.B was supported by NIH K08 HL169725. R.J.G. was additionally supported by the Robert J. Anthony Fund for Cardiovascular Research and the JB Cardiovascular Research Fund, and L.J.S. was partially supported by NIH R01 HD095881.

## AUTHOR CONTRIBUTIONS

S.-L.L. conceived and directed the project. R.J.G led the clinical study/experimental design and implementation. P.L. performed the experiments and data processing and analyses. Y.X. and K.X. performed molecular modeling and data analyses. Y.L. assisted experiments. D.J. led SARS-CoV-2 variant genotyping and DNA sequencing analyses. C.C., J.S.B., J.C.H., R.M., and R.J.G. provided clinical samples and related information. C.C.H, M.C., and J.L. provided hamster serum samples and associated information. P.L., J.N.F. and S.-L.L. wrote the paper. Y.-M.Z, L.J.S., E.M.O. provided insightful discussion and revision of the manuscript.

## DECLARATION OF INTERESTS

The authors do not declare any competing interests.

## METHODS

### Lead Contact

Dr. Shan-Lu Liu can be reached at with requests for reagents and other resources.

### Materials Availability

Contact Dr. Shan-Lu Liu with questions regarding requests for materials.

### Data and Code Availability

Our study does not report original code. Data can be requested from Dr. Shan-Lu Liu.

## EXPERIMENTAL MODEL AND SUBJECT DETAILS

### Vaccinated and patient cohorts

All data were collected from our cohorts under approved IRB protocols as follows: Bivalent mRNA vaccination: 2020H0228, 2020H0527, and 2017H0292; BA.2.86/JN.1 wave patients: 2020H0527, 2020H0531, 2020H0240, and 2020H0175; XBB.1.5 monovalent-vaccinated hamsters: 2009A1060-R4 and 2020A00000053-R1. The first human cohort was the Ohio State University Wexner Medical Center healthcare workers (HCWs) that received at least two doses of monovalent WT mRNA vaccine and a dose of the bivalent (WT + BA.4/5) mRNA booster vaccine (n=10) (**Table S1**). All individuals were administered two homologous doses of mRNA vaccine, 5 received Moderna and 5 Pfizer. Nine individuals received a third dose of vaccine (4 Moderna, 5 Pfizer) while 1 individual did not receive a third dose. Five individuals were administered the Pfizer formulation of the bivalent vaccine while 5 received the Moderna formulation. Blood was collected between 23-108 days post bivalent dose administration. Individuals ranged from 27-46 years old with a median of 37 males and 5 females were recruited.

The second cohort of human samples were patients at the Ohio State Wexner Medical center that were either admitted to the ICU during the BA.2.86/JN.1 wave of infection in Columbus, OH (11/23/2024-8/11/2024) (n=6) or collected from first responders and household contacts in the STOP-COVID cohort that were symptomatic during that time period (n=4). Positivity for SARS-CoV-2 infection was confirmed via RT-PCR and the infecting variant was determined through sequencing of nasopharyngeal swabs and next generation sequencing (Artic v5.3.2, IDT, Coralville, IA and Aritc v4.1 primers, Illumina, San Diego, CA). Ages ranged from 34-81 with a median of 52. 4 females and 6 males were recruited to this cohort.

The last cohort were golden Syrian hamsters (Envigo, Indianapolis, IN) that received recombinant mumps virus vaccines encoding monovalent XBB.1.5 spike (n=10). The vaccine was delivered intranasally at 1.5 x 10^5^ PFU twice three weeks apart. Hamsters were all 15 weeks of age and blood was collected 2 weeks after the booster dose was administered.

### Cell lines and maintenance

Human epithelial kidney cells (293T, ATCC, RRID: CVCL_1926) and 293T cells overexpressing human ACE2 (293T-ACE2) (BEI Resources, RRID: CVCL_A7UK) were maintained in DMEM (Signma Aldrich, Cat #11965-092) supplemented with 10% fetal bovine serum (Thermo Fisher, Cat #F1051) and 0.5% penicillin/streptomycin (HyClone, Cat #SV30010). Human lung adenocarcinoma cell line CaLu-3 cells were maintained in EMEM (ATCC, Cat #30-2003) supplemented with the same components. To passage, cells were washed in phosphate-buffered saline then detached using 0.05% Trypsin + 0.53 mM EDTA (Corning, Cat #27106). Cells were maintained at 37°C with 5.0% CO_2_.

## METHOD DETAILS

### Plasmids

Spike plasmids are engineered into the pcDNA3.1 plasmid backbone with a FLAG tag at C-terminal end of the coding sequence except D614G, which has a FLAG tag at both N- and C-terminal ends. D614G was synthesized and cloned into pcDNA3.1 using KpnI/BamHI restriction enzymes by GenScript Biotech. JN.1 spike was generated through site-directed mutagenesis from BA.2.86 (synthesized by GenScript) and each of the other variants were generated by site-directed mutagenesis from JN.1^8,9^. Our pseudotyped HIV-1 vectors are based on the pNL4-3-inGluc originally received from David Derse (NIH), with modifications by Marc Johnson^53^.

### Pseudotyped lentivirus production and infectivity

Pseudotyped viruses were produced via polyethyleneimine transfection (Transporter 5 Transfection Reagent, Polyscienes, Cat #26008-5) of 293T cells with a 2:1 ratio of pNL43-inGluc vector and spike^52^. Viruses were collected 48 and 72 hours post-transfection and used to infect target cells 293T-ACE2 and CaLu-3 cells. To measure these readouts, equal volumes of infected cell media and *Gaussia* luciferase substrate (0.1 M Tris pH 7.4, 0.3 M sodium ascorbate, 10 µM coelenterazine) are combined and luminescence is determined by a Cytation 5 Imaging Reader (BioTek). These readings are taken 48 and 72 hours post-infection.

### Virus neutralization assay

Viral infectivity is determined for each variant and normalized to ensure that comparable infectious viral particles were used for this assay^52^. Sera from the various cohorts was serially diluted to final dilutions 1:40, 1:160, 1:640, 1:2560, 1:10240 and one no-sera well for each individual sample. S309 was diluted to 12, 3, 0.75, 0.19, 0.047 and 0 μg/mL. Equal volumes of normalized vector were added to the serially diluted sera and incubated for 1 hour at 37°C. The mixtures were then used to infect 293T-ACE2 cells and relative infectivity determined at 48 and 72 hours post infection as described above. Neutralization titers at 50% were calculated via least squares fit nonlinear regression using GraphPad v10 (San Diego, CV) with values normalized to the no sera/antibody control.

### Antigenic cartography analysis

Racmacs v1.1.35 was used to generate the antigenic maps^54^. Briefly, instructions detailed on the GitHub entry (https://github.com/acorg/Racmacs/tree/master) were used to run the program in R (Vienna, Austria). Raw neutralization titers are input into the program where they are then log2 transformed and plotted in a distance table. This distance table is then used to perform multidimensional scaling and lot the individual sera samples (squares) and antigens (circles) in two-dimensional space. These plots are scaled by antigenic distance units (AU) where 1 AU = about a two-fold difference in nAb titer. Program optimizations were kept on default and maps were exported using the “view(map” function and labeled using Microsoft Office PowerPoint.

### Cell-cell fusion

Cell-cell fusion was performed as previously described^20^. 293T cells were co-transfected with spike plasmids and GFP. The cells were then detached using Trypsin + 0.53 mM EDTA and co-cultured with either 293T-ACE2 or CaLu-3 cells. Cells were co-cultured for 6.5 hours (293T-ACE2) or 4 hours (CaLu-3) before fusion was imaged using a Leica DMi8 fluorescence microscope. The Leica X Applications Suite was used quantify total areas of fusion by outlining areas of GFP fluorescence and calculating area within these spaces. Scale bars represent 150 µM. Three representative images were taken for each variant and used for quantification; one representative image was chosen for presentation in **Fig 5**.

### Spike surface expression

Surface expression of spike was determined on 293T cells used to produce pseudotyped viruses. After collection of virus 72 hours post-transfection, cells were detached using PBS + 5 mM EDTA and then fixed in 3.7% formaldehyde. Cells were stained with an anti-S1 polyclonal antibody (Sino Biological, T62-40591, RRID:AB_2893171) and anti-Rabbit-IgG FITC secondary antibody (Sigma, F9887, RRID:AB_259816). Flow cytometry data was collected using an Attune NxT flow cytometer and analyzed using FlowJo v10.8.1.

### Spike processing

Spike processing by furin was determined by lysing 293T cells producing pseudotyped viruses using RIPA buffer (Sigma Aldrich, R0278) plus protease inhibitor cocktails (Sigma, P8340). Samples were run on a 10% SDS-polyacrylamide gel and transferred onto a PVDF membrane. Blots were probed with anti-S2 (Sino Biological, T62-40590, RRID:AB_2857932), anti-S1 (Sino Bio, T62-40591, RRID:AB_2893171), anti-p24 (Abcam, ab63917; NIH ARP-1513), and anti-GAPDH (Proteintech, 10028230) antibodies, respectively. Secondary antibodies used were anti-Rabbit-IgG-HRP (Sigma, Cat#A9169, RRID:AB_258434) and anti-Mouse-IgG-HRP (Sigma, Cat#A5728, RRID:AB_258232). Chemiluminescence was determined by applying Immobilon Crescendo Western HRP substrate (Millipore, WBLUR0500) to the blots followed by immediately reading on a GE Amersham Imager 600. NIH ImageJ (Bethesda, MD) was used to quantify S2/S and S1/S ratios based on relative band intensity.

### S1 shedding

HEK293T cells were transfected with spike expression constructs. Twenty-four hours after transfection, cells were treated with or without sACE2 (10 μg/mL) for 4 hours at 37°C. Cell lysates and culture media were harvested. S1-containig cell culture media were incubated with 10 μL of protein A/G-conjugated anti-S1 beads (Santa Cruz, sc-2003) overnight to precipitate S1 subunit. Following immunoprecipitation, cell lysates and shed S1 were run on 10% SDS-PAGE, transferred to membranes, and probed with anti-S1 (Sino Biological, T62-40591, RRID:AB_2893171), anti-S2 (Sino Biological, T62-40590, RRID:AB_2857932) and anti-GAPDH (Proteintech, 10028230) antibodies, respectively. Anti-mouse-IgG-Peroxidase (Sigma, A5278) and anti-rabbit-IgG-HRP (Sigma, A9169) were used as secondary antibodies.

### Virus inactivation by temperature

Pseudotyped lentiviruses were pelleted through 20% sucrose in TMS buffer (25 mM Tris, 25 mM maleic acid, 150 mM NaCl, pH 6.5) by centrifugation at 25,000 g at 4°C for 2 h in Beckman SW41 rotor. Viruses were resuspended in DMEM (pH 7.4) without serum, incubated at different temperatures (37 to 43°C) for 1 h, and inoculated onto 293T-ACE2 cells to assay the transduction efficiency. Viruses stayed at 4° C throughout the treatment served as control for comparison.

### Structural modeling and analyses

Structural modeling to assess the impact of spike mutations on ACE2 binding, conformational stability, and antibody evasion was performed using the SWISS-MODEL server. Glycosylation modifications at residues N30 and N61 were incorporated using the program Coot. This analysis utilized published X-ray crystallography and cryo-EM structures (PDB: 8X4H, 8Y5J, 6LZG, 7XEG, 7KMG, 7YAD, 8DLS, 7UAP, 7UAR) as templates. The potential effects of key mutations on these interactions were examined, and the resulting models were visually represented using PyMOL.

### Quantification and statistical analysis

All statistical analyses in this work were conducted using GraphPad Prism 10. NT_50_ values were calculated by least-squares fit non-linear regression. Error bars in Figures 1D, 1E, 5B, 5D and 6B represent means ± standard errors. Error bars in Figures 2A, 2C, and 2E represent geometric means with 95% confidence intervals. Error bars in Figure 3A represent means ± standard deviation. Statistical significance was analyzed using log10 transformed NT_50_ values to better approximate normality (Figures 2A, 2C and 2E), and multiple groups comparisons were made using a one-way ANOVA with Bonferroni post-test. Cell-cell fusion was quantified using the Leica X Applications Suite software (Figures 5A and 5C). S processing was quantified by NIH ImageJ (Figure 6C and 7C).

## Figure Legends

**Figure S1** Antigenic distances of JN.1-derived subvariants relative to D614G or JN.1 in three groups of cohorts. **(A)** bivalent-vaccinated HCWs; **(B)** BA.2.86/JN.1-wave infected people; and **(C)** XBB.1.5-vaccinated hamsters. One antigenic distance unit (AU) is equivalent to a 2-fold difference in nAb titer shown in Figure 2.

**Figure S2** NAb titers in the sera of **(A)** first responders and household contacts during the BA.2.86/JN.1 in Columbus who became COVID positive and suffered mild illness (n = 4) and **(B)** ICU patients during the BA.2.86/JN.1-wave in Columbus, Ohio (n=6).

## Notes

### Competing Interest Statement

The authors have declared no competing interest.

## REFERENCES

1 CDC. COVID Data Tracker, Variant Proportions (2024).

2 Gangavarapu, K. et al. Outbreak.info genomic reports: scalable and dynamic surveillance of SARS-CoV-2 variants and mutations. Nat Methods 20, 512–522, doi:10.1038/s41592-023-01769-3 (2023).

3 Yang, S. et al. Fast evolution of SARS-CoV-2 BA.2.86 to JN.1 under heavy immune pressure. The Lancet. Infectious Diseases 24, e70–e72, doi:10.1016/S1473-3099(23)00744-2 (2024).

4 Wang, X., Lu, L. & Jiang, S. SARS-CoV-2 evolution from the BA.2.86 to JN.1 variants: unexpected consequences. Trends in Immunology 45, 81–84, doi:10.1016/j.it.2024.01.003 (2024).

5 Planas, D. et al. Distinct evolution of SARS-CoV-2 Omicron XBB and BA.2.86/JN.1 lineages combining increased fitness and antibody evasion. Nature Communications 15, 2254, doi:10.1038/s41467-024-46490-7 (2024).

6 Kaku, Y. et al. Virological characteristics of the SARS-CoV-2 JN.1 variant. The Lancet. Infectious Diseases 24, e82, doi:10.1016/S1473-3099(23)00813-7 (2024).

7 Wang, Q. et al. XBB.1.5 monovalent mRNA vaccine booster elicits robust neutralizing antibodies against XBB subvariants and JN.1. Cell Host & Microbe 32, 315–321 e313, doi:10.1016/j.chom.2024.01.014 (2024).

8 Qu, P. et al. Immune evasion, infectivity, and fusogenicity of SARS-CoV-2 BA.2.86 and FLip variants. Cell 187, 585–595 e586, doi:10.1016/j.cell.2023.12.026 (2024).

9 Li, P. et al. Neutralization escape, infectivity, and membrane fusion of JN.1-derived SARS-CoV-2 SLip, FLiRT, and KP.2 variants. Cell Reports 43, 114520, doi:10.1016/j.celrep.2024.114520 (2024).

10 Jain, S. et al. XBB.1.5 monovalent booster improves antibody binding and neutralization against emerging SARS-CoV-2 Omicron variants. bioRxiv, 2024.2002.2003.578771, doi:10.1101/2024.02.03.578771 (2024).

11 Wang, X., Jiang, S., Ma, W., Zhang, Y. & Wang, P. Robust neutralization of SARS-CoV-2 variants including JN.1 and BA.2.87.1 by trivalent XBB vaccine-induced antibodies. Signal Transduction and Targeted Therapy 9, 123, doi:10.1038/s41392-024-01849-6 (2024).

12 Lasrado, N., Rössler, A., Rowe, M., Collier, A. Y. & Barouch, D. H. Neutralization of SARS-CoV-2 Omicron subvariant BA.2.87.1. Vaccine 42, 2117–2121, doi:10.1016/j.vaccine.2024.03.007 (2024).

13 Murrell, B. SARS-CoV-2 Lineage Competition (2024-06-13), <https://github.com/MurrellGroup/lineages> (2024).

14 Li, P. et al. Distinct patterns of SARS-CoV-2 BA.2.87.1 and JN.1 variants in immune evasion, antigenicity, and cell-cell fusion. mBio 15, e0075124, doi:10.1128/mbio.00751-24 (2024).

15 Zhang, L. et al. Rapid spread of the SARS-CoV-2 JN.1 lineage is associated with increased neutralization evasion. iScience 27, 109904, doi:10.1016/j.isci.2024.109904 (2024).

16 Gonzalez-Reiche, A. S. et al. Sequential intrahost evolution and onward transmission of SARS-CoV-2 variants. Nature Communications 14, 3235, doi:10.1038/s41467-023-38867-x (2023).

17 Dadonaite, B. et al. Spike deep mutational scanning helps predict success of SARS-CoV-2 clades. Nature 631, 617–626, doi:10.1038/s41586-024-07636-1 (2024).

18 Kaku, Y. et al. Virological characteristics of the SARS-CoV-2 KP.2 variant. The Lancet. Infectious Diseases 24, e416, doi:10.1016/s1473-3099(24)00298-6 (2024).

19 Jian, F. et al. Convergent evolution of SARS-CoV-2 XBB lineages on receptor-binding domain 455-456 synergistically enhances antibody evasion and ACE2 binding. PLoS Pathog 19, e1011868, doi:10.1371/journal.ppat.1011868 (2023).

20 Qu, P. et al. Evasion of neutralizing antibody responses by the SARS-CoV-2 BA.2.75 variant. Cell Host & Microbe 30, 1518–1526 e1514, doi:10.1016/j.chom.2022.09.015 (2022).

21 Qu, P. et al. Enhanced evasion of neutralizing antibody response by Omicron XBB.1.5, CH.1.1, and CA.3.1 variants. Cell Reports 42, 112443, doi:10.1016/j.celrep.2023.112443 (2023).

22 Cao, Y. et al. Imprinted SARS-CoV-2 humoral immunity induces convergent Omicron RBD evolution. Nature 614, 521–529, doi:10.1038/s41586-022-05644-7 (2023).

23 Wang, Q. et al. Alarming antibody evasion properties of rising SARS-CoV-2 BQ and XBB subvariants. Cell 186, 279–286.e278, doi:10.1016/j.cell.2022.12.018 (2023).

24 Zhou, T. et al. Structural basis for potent antibody neutralization of SARS-CoV-2 variants including B.1.1.529. Science (New York, N.Y.) 376, eabn8897, doi:10.1126/science.abn8897 (2022).

25 Pinto, D. et al. Cross-neutralization of SARS-CoV-2 by a human monoclonal SARS-CoV antibody. Nature 583, 290–295, doi:10.1038/s41586-020-2349-y (2020).

26 Evans, J. P. et al. Neutralization of SARS-CoV-2 Omicron sub-lineages BA.1, BA.1.1, and BA.2. Cell Host & Microbe 30, 1093–1102 e1093, doi:10.1016/j.chom.2022.04.014 (2022).

27 Arora, P. et al. Lung cell entry, cell-cell fusion capacity, and neutralisation sensitivity of omicron sublineage BA.2.75. The Lancet. Infectious Diseases 22, 1537–1538, doi:10.1016/s1473-3099(22)00591-6 (2022).

28 Wang, X. et al. Enhanced neutralization of SARS-CoV-2 variant BA.2.86 and XBB sub-lineages by a tetravalent COVID-19 vaccine booster. Cell Host & Microbe 32, 25–34.e25, doi:10.1016/j.chom.2023.11.012 (2024).

29 Barouch, D. H. Covid-19 Vaccines - Immunity, Variants, Boosters. The New England Journal of Medicine 387, 1011–1020, doi:10.1056/NEJMra2206573 (2022).

30 Kaku, Y., Uriu, K., Okumura, K., Ito, J. & Sato, K. Virological characteristics of the SARS-CoV-2 KP.3.1.1 variant. The Lancet. Infectious diseases, doi:10.1016/s1473-3099(24)00505-x (2024).

31 Jian, F. et al. Evolving antibody response to SARS-CoV-2 antigenic shift from XBB to JN.1. bioRxiv, 2024.2004.2019.590276, doi:10.1101/2024.04.19.590276 (2024).

32 Kaku, Y. et al. Virological characteristics of the SARS-CoV-2 KP.3, LB.1, and KP.2.3 variants. The Lancet. Infectious Diseases 24, e482–e483, doi:10.1016/S1473-3099(24)00415-8 (2024).

33 Faraone, J. N. & Liu, S. L. Immune imprinting as a barrier to effective COVID-19 vaccines. *Cell reports*. Medicine 4, 101291, doi:10.1016/j.xcrm.2023.101291 (2023).

34 Evans, J. P. & Liu, S. L. Challenges and Prospects in Developing Future SARS-CoV-2 Vaccines: Overcoming Original Antigenic Sin and Inducing Broadly Neutralizing Antibodies. Journal of Immunology 211, 1459–1467, doi:10.4049/jimmunol.2300315 (2023).

35 San Filippo, S., et al. Comparative Efficacy of Early COVID-19 Monoclonal Antibody Therapies: A Retrospective Analysis. Open Forum Infectious Diseases 9, ofac080, doi:10.1093/ofid/ofac080 (2022).

36 Liu, Z. et al. Neutralization of SARS-CoV-2 BA.2.86 and JN.1 by CF501 adjuvant-enhanced immune responses targeting the conserved epitopes in ancestral RBD. Cell Reports. Medicine 5, 101445, doi:10.1016/j.xcrm.2024.101445 (2024).

37 Faraone, J. N. et al. Neutralization escape of Omicron XBB, BR.2, and BA.2.3.20 subvariants. Cell Reports. Medicine 4, 101049, doi:10.1016/j.xcrm.2023.101049 (2023).

38 Faraone, J. N. et al. Immune evasion and membrane fusion of SARS-CoV-2 XBB subvariants EG.5.1 and XBB.2.3. Emerg Microbes Infect 12, 2270069, doi:10.1080/22221751.2023.2270069 (2023).

39 Chen, Y. et al. Broadly neutralizing antibodies to SARS-CoV-2 and other human coronaviruses. Nature Reviews Immunology 23, 189–199, doi:10.1038/s41577-022-00784-3 (2023).

40 Zhang, L. et al. SARS-CoV-2 BA.2.86 enters lung cells and evades neutralizing antibodies with high efficiency. Cell 187, 596–608.e517, doi:10.1016/j.cell.2023.12.025 (2024).

41 Cai, Y. et al. AZD3152 neutralizes SARS-CoV-2 historical and contemporary variants and is protective in hamsters and well tolerated in adults. Sci Transl Med 16, eado2817, doi:10.1126/scitranslmed.ado2817 (2024).

42 Francica, J. R., et al. 1355. The SARS-CoV-2 Monoclonal Antibody AZD3152 Potently Neutralizes Historical and Emerging Variants and is Being Developed for the Prevention and Treatment of COVID-19 in High-risk Individuals. Open Forum Infect Dis. 2023 Nov 27;10(Suppl 2):ofad500.1192. doi: 10.1093/ofid/ofad500.1192. eCollection (2023).

43 Cao, Y. et al. Rational identification of potent and broad sarbecovirus-neutralizing antibody cocktails from SARS convalescents. Cell Reports 41, 111845, doi:10.1016/j.celrep.2022.111845 (2022).

44 Wang, Q., Guo, Y., Ho, J. & Ho, D. D. Pemivibart is less active against recent SARS-CoV-2 JN.1 sublineages. bioRxiv, 2024.2008.2012.607496, doi:10.1101/2024.08.12.607496 (2024).

45 Planas, D. et al. Escape of SARS-CoV-2 variants KP1.1, LB.1 and KP3.3 from approved monoclonal antibodies. bioRxiv, 2024.2008.2020.608835, doi:10.1101/2024.08.20.608835 (2024).

46 Benton, D. J. et al. Receptor binding and priming of the spike protein of SARS-CoV-2 for membrane fusion. Nature 588, 327–330, doi:10.1038/s41586-020-2772-0 (2020).

47 Zhang, S. et al. Loss of Spike N370 glycosylation as an important evolutionary event for the enhanced infectivity of SARS-CoV-2. Cell Research 32, 315–318, doi:10.1038/s41422-021-00600-y (2022).

48 Raisinghani, N., Alshahrani, M., Gupta, G. & Verkhivker, G. Atomistic Prediction of Structures, Conformational Ensembles and Binding Energetics for the SARS-CoV-2 Spike JN.1, KP.2 and KP.3 Variants Using AlphaFold2 and Molecular Dynamics Simulations: Mutational Profiling and Binding Free Energy Analysis Reveal Epistatic Hotspots of the ACE2 Affinity and Immune Escape. bioRxiv, 2024.2007.2009.602810, doi:10.1101/2024.07.09.602810 (2024).

49 Taylor, A. L. & Starr, T. N. Deep mutational scanning of SARS-CoV-2 Omicron BA.2.86 and epistatic emergence of the KP.3 variant. bioRxiv, doi:10.1101/2024.07.23.604853 (2024).

50 Zeng, C. et al. Neutralization and Stability of SARS-CoV-2 Omicron Variant. bioRxiv, 2021.2012.2016.472934,doi:10.1101/2021.12.16.472934 (2021).

51 FDA. Updated COVID-19 Vaccines for Use in the United States Beginning in Fall 2024 (2024).

52 Zeng, C. et al. Neutralizing antibody against SARS-CoV-2 spike in COVID-19 patients, health care workers, and convalescent plasma donors. JCI Insight 5, doi:10.1172/jci.insight.143213 (2020).

53 Mazurov, D., Ilinskaya, A., Heidecker, G., Lloyd, P. & Derse, D. Quantitative comparison of HTLV-1 and HIV-1 cell-to-cell infection with new replication dependent vectors. PLoS Pathog 6, e1000788, doi:10.1371/journal.ppat.1000788 (2010).

54 Smith, D. J. et al. Mapping the antigenic and genetic evolution of influenza virus. Science 305, 371–376, doi:10.1126/science.1097211 (2004).

