## Supplemental Figures for "Neutralization and Stability of JN.1-derived LB.1, KP.2.3, KP.3 and KP.3.1.1 Subvariants"

**Table S1. Bivalent-vaccinated healthcare workers (HCWs) and BA.2.86/JN.1-wave patients**

| | **Bivalent Health Care Workers** (n=10) | **BA.2.86/JN.1 Wave Patients** (n=10) |
|---|---|---|
| **Age in Years at Sample Collection [Median (Range)]** | 37 (27-46) | 52 (34-81) |
| **Gender** [n (% of Total)] | | |
| Male | 5 (50%) | 6 (60%) |
| Female | 5 (50%) | 4 (40%) |
| **Sample Collection Window** | Dec. 2022- Jan.2023 | Nov. 2023-Aug.2024 |
| **Vaccine status** [n (% of Total)] | NA | |
| 1-dose Pfizer | NA | 1(10%) |
| 2-dose Moderna | NA | 2 (20%) |
| 3-dose Moderna | NA | 1 (10%) |
| 4-dose Moderna | NA | 1 (10%) |
| 1-dose Moderna +1-dose Pfizer bivalent | NA | 1 (10%) |
| 1-dose Pfizer +1-dose Pfizer bivalent | NA | 2 (20%) |
| 2-dose Pfizer +1-dose Pfizer bivalent | 1 (10%) | NA |
| 3-dose Pfizer +1-dose Moderna bivalent | NA | 1 (14.3%) |
| 3-dose Pfizer +1-dose Pfizer bivalent | 3 (30%) | NA |
| 3-dose Pfizer +1-dose Moderna | 1 (10%) | NA |
| 3-dose Moderna +1-dose Moderna bivalent | 4 (40%) | 1 (14.3%) |
| 2-dose Moderna +1 Pfizer +1-dose Pfizer bivalent | 1 (10%) | NA |
| Days from last vaccination | NA | 675 (34-1033) |
| Days post the bivalent dose for recipients | 66 (23-108) | NA |
| **COVID-19 positive** [n (% of Total)] | 8 (80%) | 10 (100%) |
| Days before sample collection [(Median Range)] | 276.5 (182-994) | 7 (1-10) |
| **Infected Variants** | | |
| JN.1/BA.2.86 | NA | 2 (20%) |
| Undetermined | NA | 8 (80%) |

Summary of the demographic information for two cohorts used for neutralization experiments depicted in Figure 2. “NA” means the category is not applicable to the cohort.

A

| Bivalent HCWs | | | |
|---|---|---|---|
| AD (D614G) | | AD (JN.1) | |
| JN.1 | 3.2 | D614G | 3.2 |
| FLiRT | 4.2 | FLiRT | 1.0 |
| FLiRT_DelS31 | 6.4 | FLiRT_DelS31 | 5.4 |
| FLiRT_Q183H | 3.9 | FLiRT_Q183H | 0.7 |
| LB.1 | 6.4 | LB.1 | 6.0 |
| KP.2 | 3.7 | KP.2 | 1.9 |
| KP.2_DelS31 | 6.1 | KP.2_DelS31 | 4.7 |
| KP.2_H146Q | 3.9 | KP.2_H146Q | 2.4 |
| KP.2.3 | 6.4 | KP.2.3 | 5.6 |
| KP.3 | 4.0 | KP.3 | 1.6 |
| KP.3.1.1 | 6.5 | KP.3.1.1 | 5.3 |

B

| BA.2.86/JN.1-wave patients | | | |
|---|---|---|---|
| AD (D614G) | | AD (JN.1) | |
| JN.1 | 2.3 | D614G | 2.3 |
| FLiRT | 2.7 | FLiRT | 1.0 |
| FLiRT_DelS31 | 5.0 | FLiRT_DelS31 | 4.0 |
| FLiRT_Q183H | 2.7 | FLiRT_Q183H | 1.1 |
| LB.1 | 5.3 | LB.1 | 4.5 |
| KP.2 | 2.4 | KP.2 | 0.9 |
| KP.2_DelS31 | 4.8 | KP.2_DelS31 | 3.9 |
| KP.2_H146Q | 2.4 | KP.2_H146Q | 0.8 |
| KP.2.3 | 4.7 | KP.2.3 | 3.0 |
| KP.3 | 3.2 | KP.3 | 1.5 |
| KP.3.1.1 | 4.8 | KP.3.1.1 | 4.6 |

C

| XBB.1.5-monovalent hamsters | | | |
|---|---|---|---|
| AD (D614G) | | AD (JN.1) | |
| JN.1 | 1.7 | D614G | 1.6 |
| FLiRT | 1.8 | FLiRT | 0.4 |
| FLiRT_DelS31 | 2.9 | FLiRT_DelS31 | 1.9 |
| FLiRT_Q183H | 1.6 | FLiRT_Q183H | 0.4 |
| LB.1 | 2.8 | LB.1 | 1.8 |
| KP.2 | 1.9 | KP.2 | 0.3 |
| KP.2_DelS31 | 2.8 | KP.2_DelS31 | 1.5 |
| KP.2_H146Q | 2.1 | KP.2_H146Q | 0.5 |
| KP.2.3 | 3.0 | KP.2.3 | 1.8 |
| KP.3 | 1.9 | KP.3 | 0.3 |
| KP.3.1.1 | 2.8 | KP.3.1.1 | 1.5 |

**Figure S1**

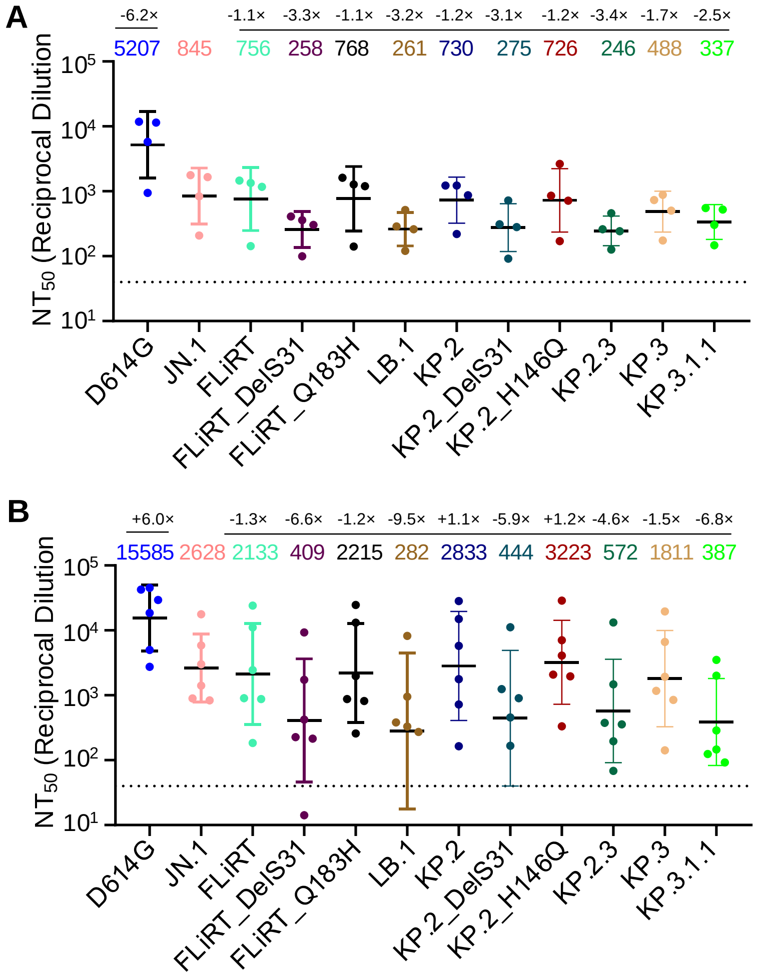
A
-6.2×
-1.1× -3.3× -1.1× -3.2× -1.2× -3.1× -1.2× -3.4× -1.7× -2.5×
5207 845 756 258 768 261 730 275 726 246 488 337
NT50 (Reciprocal Dilution
D614G
JN.1
FLiRT
FLiRT_DelS31
FLiRT_Q183H
LB.1
KP.2
KP.2_DelS31
KP.2_H146Q
KP.2.3
KP.3
KP.3.1.1
B
+6.0×
-1.3× -6.6× -1.2× -9.5× +1.1× -5.9× +1.2× -4.6× -1.5× -6.8×
15585 2628 2133 409 2215 282 2833 444 3223 572 1811 387

**Figure S2**
